# The transcription factor Vca0578 (DsvR) mediated expression of ZapC is required to promote cell division during lytic transglycosylase insufficiency in *Vibrio cholerae*

**DOI:** 10.64898/2026.04.01.715812

**Authors:** Upasana Basu, Anna Isabell Weaver, Nicole Lin, Asraa Ahmed, Sebastian Krautwurst, Kai Papenfort, Tobias Dörr

**Affiliations:** Weill Institute for Cell and Molecular Biology, Cornell University, Ithaca, United States; College of Veterinary Medicine, Cornell University, Ithaca, United States; Institute of Microbiology, Faculty of Biological Sciences, Friedrich Schiller University, Jena, Germany; Microverse Cluster, Friedrich Schiller University, Jena, Germany; Department of Microbiology, Cornell University, Ithaca, United States; Cornell Institute for Host-Microbe Interactions and Disease (CIHMID), Ithaca, United States

**Keywords:** Peptidoglycan, Lytic transglycosylase, Cell division, Zap protein, Z-ring

## Abstract

The bacterial peptidoglycan (PG) cell wall, a polymer made of amino-acid-bearing glycan strands, maintains cell shape, provides structural integrity, and protects against osmotic lysis. PG maintenance is an active process that requires regulated PG breakdown to make space for insertion of new PG strands. PG breakdown is accomplished by ‘autolysins’, i.e. endogenous enzymes with cell wall cleavage activity. The lytic transglycosylases (LTGs), a class of autolysins, for example, cleave glycan strands during PG remodelling. LTGs are broadly conserved and are highly redundant in bacteria, but their physiological role is poorly-defined. In this study, we interrogated physiological consequences of LTG insufficiency in *Vibrio cholerae* using TnSeq to gain insights about roles of these enzymes. We identify an uncharacterized transcription factor, Vca0578, which alleviates defects associated with the Δ6LTG mutant. We demonstrate that Vca0578 positively regulates the expression of *zapC*, a typically non-essential Z-ring associated protein. In the absence of *zapC*, cell division was impaired during perturbations of cell envelope homeostasis caused by absence of LTGs, or by exposure to antibiotics inhibiting cell elongation; either condition rendered *zapC* conditionally essential. This essentiality could be overcome by increasing FtsZ levels. Lastly, we found that ZapC also contributes to both width and length homeostasis during normal growth. This work thus uncovers a novel transcriptional circuit that contributes to effective cell division in *Δ*6LTG cells, and suggests an essential role for ZapC in cell division under stress conditions that cause perturbation of cell width homeostasis.

**AUTHOR SUMMARY:** Bacteria must maintain their outer shell (the cell envelope) in the face of changes in the environment. For this, they use elaborate systems that remodel the cell envelope. How some of these systems work is not well understood. In this study, we describe a new gene circuit that is required to keep cells dividing when the cell envelope is compromised. We found that Vca0578, a putative transcription factor, controls expression of the *zapC* gene. The protein ZapC then helps bacteria grow and divide when the cell envelope is under stress, for example, in the presence of certain antibiotics. Thus, we have discovered a regulatory circuit that promotes bacterial growth and antibiotic resistance under stress.

## INTRODUCTION

Most bacteria are surrounded by a peptidoglycan (PG) cell wall that protects them against lysis from internal turgor pressure and external environmental challenges ^1–3^. PG resides in the periplasm, the space between the inner membrane (IM) and the outer membrane (OM) in Gram-negative bacteria. This sacculus-like structure is made of glycan strands that are crosslinked to one another to form one continuous and covalently closed macromolecule. Glycosyltransferases (GTases) (aPBPs or SEDS-proteins) catalyze formation of β-1,4 glycosidic linkages between heterodimers consisting of alternating N-acetylglucosamine (GlcNAc) and N-acetylmuramic acid (MurNAc) residues ^4^. Each MurNAc has a pentapeptide attachment consisting of ^1^L-alanine - ^2^D-glutamate - ^3^meso-diaminopimelic acid (DAP) - ^4^D-alanine - ^5^D-alanine, and about 40% of these peptides are crosslinked to adjacent ones. The majority of these crosslinks are formed between an ^3^m-DAP and an adjacent ^4^D-ala resulting in the formation of D,D- or 4,3-crosslink. This reaction is catalyzed by the transpeptidase (TPase) activity of aPBPs or bPBPs. However a small fraction of L,D- or 3,3-crosslinks are formed between ^3^m-DAP residues by L,D-transpeptidases (LDTases). PG synthesis starts in the cytoplasm with the formation of lipid II, a disaccharide-pentapeptide molecule modified with an undecaprenol tail that gets flipped across the IM by the flippase MurJ. Lipid II then gets polymerized into glycan strands and crosslinked to other residues in the periplasm ^5–7^.

Growth of rod-shaped bacteria can be divided into two phases: elongation to lengthen cells, followed by division into daughter cells. Each of these processes requires active PG synthesis and remodelling at specific sites within the cell. In rod-shaped bacteria, elongation is facilitated by the elongasome machinery (also called rod-complex) to insert new PG material at scattered sites throughout the lateral cylinder. Following elongation, the divisome assembles at the mid-cell, and matures to promote septal PG synthesis (sPG). The process starts with recruitment of the bacterial tubulin homologue FtsZ to the membrane by FtsA and/or ZipA, and its polymerization to form a dynamic, condensed yet discontinuous ring-like structure called the Z-ring ^8–10^. The condensation and stability of the ring is mediated by FtsZ treadmilling dynamics ^11,12^, feedback from the sPG synthesis ^13^, and crosslinking of the FtsZ filaments by accessory ‘Zap’, the Fts**Z**-**a**ssoiated **p**roteins ^14–16^. One such accessory Zap-protein is ZapC, which is widely conserved in gammaproteobacteria and whose *in vivo* role remains elusive ^16,17^. The last essential protein to get recruited is FtsN, a bitopic membrane protein ^18,19^ that activates sPG synthesis and coordinates it with degradation ^20–24^.

Bacterial growth, that is, net expansion of the PG sacculus, requires two seemingly opposing processes: for PG to expand, degradation of existing bonds makes space for insertion of newly synthesized glycan strands ^25–30^. This action is accomplished by several classes of enzymes, collectively referred to as ‘autolysins’, which catalyze cleavage of distinct glycosidic and amide bonds of the sacculus ^31,32^. Every generation, almost half of the cell wall components are released from the sacculus, most of which is imported back into the cell through a process referred to as PG recycling ^33–35^.

Lytic transglycosylases (LTGs) are a major class of autolysins that catalyze the non-hydrolytic cleavage of the β-1,4 glycosidic linkage between MurNAc and GlcNAc residues within a glycan strand, and the mechanism for this cleavage is an intramolecular cyclization of NAM resulting in the formation of 1,6-anhydroMurNAc residue ^36^. LTGs are an enigmatic class of enzymes that have been well-characterized biochemically, but their physiological roles and regulation remains poorly understood ^37–39^. In bacteria, LTGs are very well-conserved and highly redundant, a trait usually indicative of a collectively indispensable function. For example, *Vibrio cholerae* encodes eight LTGs: MltA, MltB, MltC, MltD, MltF, RlpA (all OM-bound), MltG (IM-bound), and Slt70 (soluble). Under standard laboratory growth conditions, LTGs seem to be functionally redundant, making it challenging to associate LTGs with specific physiological roles as single-gene knockout rarely yield a strong detectable phenotype. Previous work from our lab has shown that not all LTGs are functionally equivalent, and that LTG activity is collectively essential in *V. cholerae* ^40^. Insufficient LTG activity in a minimal Δ6LTG (six out of the eight LTGs deleted, RlpA^+^ MltG^+^) strain leads to an increase in cell length. In *V. cholerae*, RlpA and MltC localize to the septum along with the sole amidase to accomplish septum cleavage and separation ^41^. Specific LTGs have been implicated in mediating cellular growth and daughter cell separation in other bacteria ^42–44^. Whether LTGs directly or indirectly contribute to cell division and elongation is subject to active inquiry. Additionally, LTGs have been implicated in terminating newly synthesized PG strands, space-makers for insertion of new strands in the sacculus, PG recycling, mitigating periplasmic crowding, and insertion of transmembrane complexes ^34,40,45–48^.

In this study we uncovered a novel transcriptional circuit, where the uncharacterized transcription factor Vca0578, here renamed to DsvR (SgrR-like **d**ivision under **s**tress **r**egulator in ***V****ibrio cholerae*), controls expression of ZapC (a non-essential Z-ring associated protein), promoting proper cell division in Δ6LTG cells. Inactivation of this pathway led to dramatic filamentation of Δ6LTG cells, suggesting that this pathway promotes cell division and length-width homeostasis during LTG insufficiency. Consistent with a role in sustaining cell division during envelope disturbances, *zapC* is also essential for intrinsic resistance to antibiotics inhibiting the rod-complex. Thus, the DsvR-ZapC pathway enables cell division under diverse cell envelope stress conditions, including that imposed by absence of multiple LTGs.

## RESULTS

### A transposon screen reveals a genetic relationship between LTGs and an uncharacterized transcription factor

A Δ6LTG *V. cholerae* strain deleted in 6 out of its 8 lytic transglycosylases (LTGs) is viable and exhibits only a minor increase in average cell length, indicating a mild cell division defect in the absence of optimal LTG activity (Fig. 1A, B). We used this strain as a tool for determining genetic interactions associated with LTG deficiency, in order to gain new insight into LTG biology. To this end, we conducted a TnSeq screen to identify genes that either mitigate or exacerbate defects in Δ6LTG, which relies on its two remaining LTGs (MltG and RlpA) for viability. This screen revealed several genes that were either conditionally essential or toxic during LTG insufficiency (Fig. 1C). A significant reduction the two remaining LTGs) answered the screen as synthetic lethal, supporting our previous results with LTG depletion strains^40^. Genes associated with outer membrane homeostasis (like O-antigen biosynthesis genes), PG recycling (like *ampG, ldcV*), and PG synthesis (like *pbp1a, lpoA*) were likewise scored as synthetic sick/lethal with Δ6LTG, perhaps indicating moderate cell envelope defects in this strain, which are exacerbated by additionally inactivating cell envelope maintenance functions. Interestingly, we also noted some genes associated with cell division (specifically the entire *tol-pal* system and *zapC*) as synthetic sick/lethal with Δ6LTG. Lastly, the screen revealed that the gene for an uncharacterized transcription factor DsvR (encoded by the gene *vca0578*) that was among the top hits from the list of conditionally essential genes (Fig. 1C). DsvR is a predicted homolog of *E. coli* SgrR, which responds to glucose-phosphate stress^49,50^, but is uncharacterized in *V. cholerae*. We thus chose to focus on the DsvR pathway for further study.

**Figure 1:**
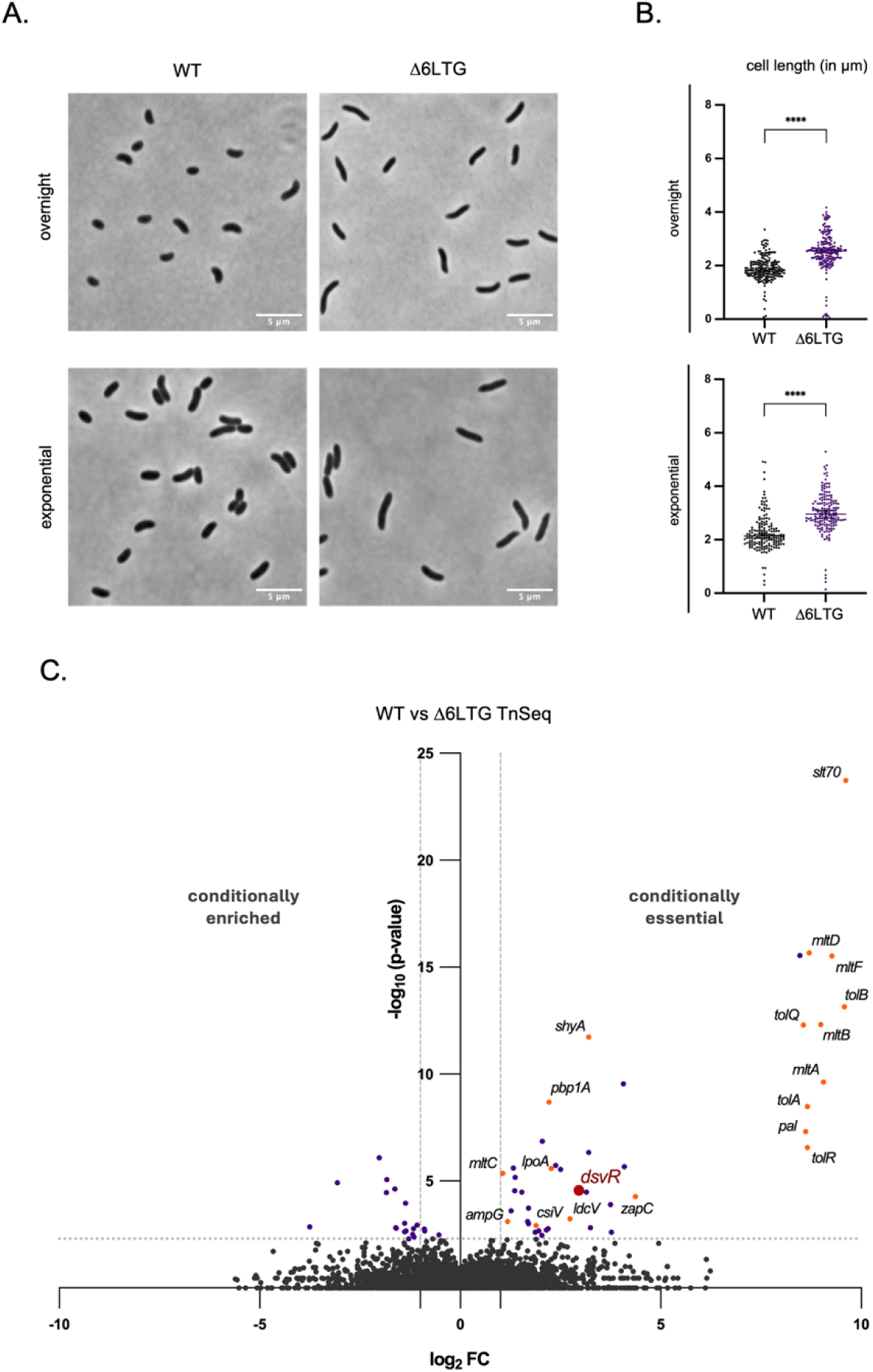
Comparison of the morphology of WT *V. cholerae* and the minimal LTG strain Δ6LTG reveal an increase in cell length and area during LTG insufficiency. (A) Overnight cultures were 1:100 sub-cultured in LB and grown at 37°C they reached the exponential stage and then imaged using phase contrast microscopy. Scale bar: 5µm; (B) Scatter plots showing length and area distributions of cells. These were analyzed using the MicrobeJ plugin in ImageJ and statistical significance was determined with Welch’s two-sample t-test. ***, p < 0.001; ****, p < 0.0001; (C) Volcano plot depicting the p-value (Y-axis) vs. the log_2_fold-change of the transposon insertion (X-axis) between WT, the control strain and Δ6LTG, the experimental strain. The dashed lines indicate the cutoff (fold-change > 2 or fold-change < −2, p-value < 0.05). An increase in fold-change indicates genes that are conditionally essential in Δ6LTG. in read-counts was observed for the 6 deleted LTGs, thus serving as internal controls for the TnSeq. Additionally, both *shyA* (encoding a major endopeptidase) and *rlpA* (one of

### The *dsvR* gene is required for proper cell growth and morphogenesis in *Δ*6LTG

To validate *dsvR* as synthetic lethal with Δ6LTG, we placed the gene under an anhydrotetracycline (aTc)-inducible promoter and grew the resulting strain in the presence or absence of aTc to observe effects of *dsvR* depletion in WT or Δ6LTG backgrounds. In the absence of aTc, we observed a >100-fold plating defect when *dsvR* was depleted in Δ6LTG background on solid agar plates (Fig. 2A). The defect was recapitulated in liquid medium, where we observed a significant reduction in growth upon *dsvR* depletion in Δ6LTG at the 6hr timepoint (Fig. 2B). We did, however, still observe residual growth upon *dsvR* depletion, indicating synthetic sickness, rather than full lethality. Consistent with this, we were able to delete the *dsvR* gene in the Δ6LTG strain. The Δ*dsvR* Δ6LTG double mutant exhibited a drastic increase in cell length, concomitant with visible lysis when plated on β-galactosidase substrate chlorophenol red-β-d-galactopyranoside (CPRG) (Fig. 3A, 3B, 3C). The morphological defects could be fully complemented upon ectopic overexpression of *dsvR* from an IPTG-inducible promoter, excluding polar effects (Fig. 3A, 3B). To characterize this mutant further, we conducted growth analysis in media with different osmolarities. The Δ*dsvR* Δ6LTG mutant exhibited significant growth defects in LB, which were exacerbated in both low- and high-salt conditions (Fig. S1), indicative of a general cell envelope defect. We then sought to determine whether Δ*dsvR* Δ6LTG formed filaments or cell chains. To this end, we generated a strain that expressed GFP constitutively to visualize the cytoplasm, and concomitantly visualized FtsZ tagged with YFP. The Δ*dsvR* Δ6LTG filaments had a causes dramatic cell division defects upstream of septation in LTG deficient cells.

**Figure 2:**
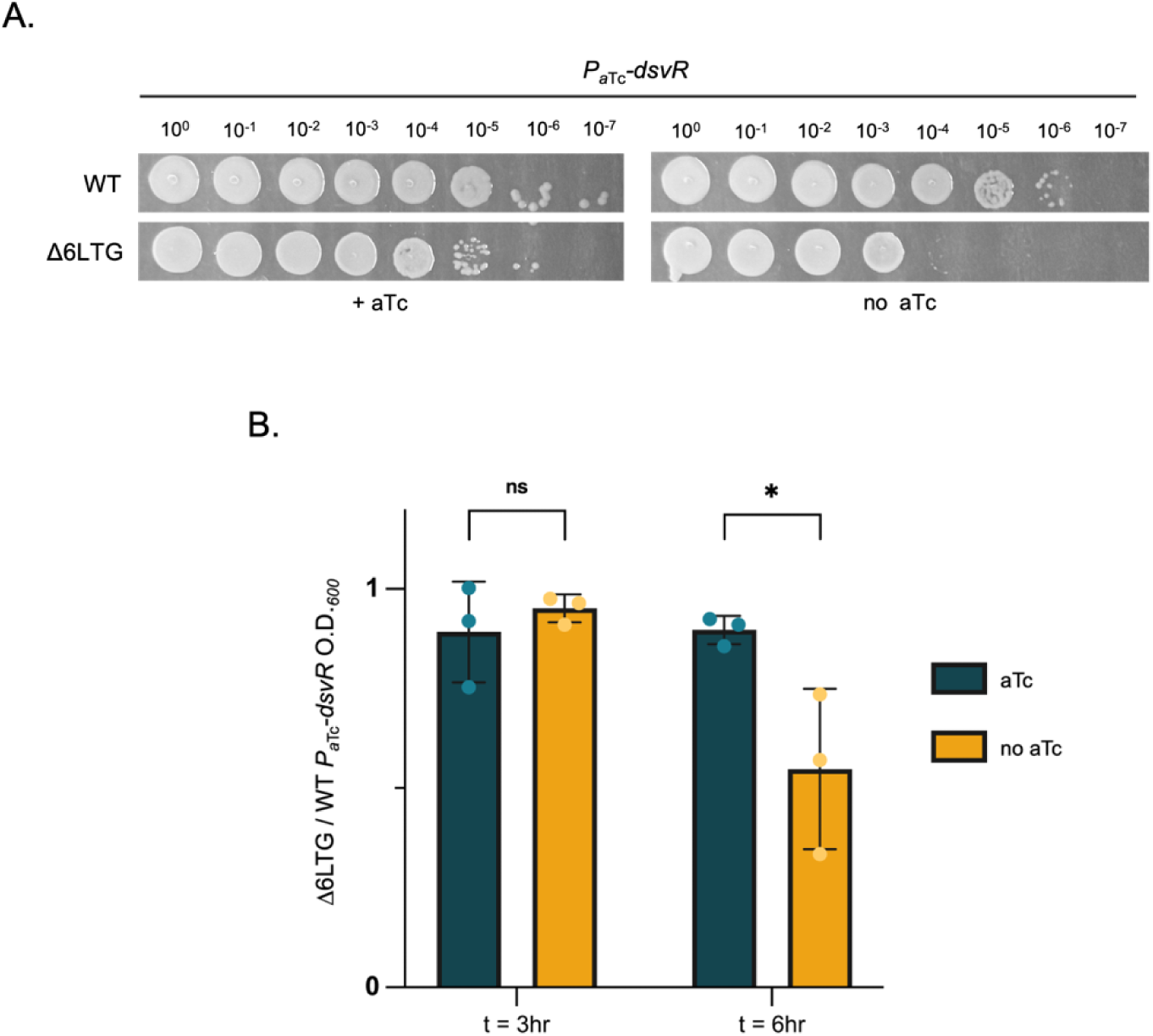
Depletion of *dsvR* in Δ6LTG leads to decreased viability. The gene *dsvR* was depleted from WT and Δ6LTG backgrounds by replacing its native promoter with an anhydrotetracycline(aTc)-inducible promoter. (A) Overnight cultures were grown in LB + aTc + chloramphenicol, and next day, they were serially diluted from 10^0^ to 10^−7^ in LB. 5µL was spotted on LB-agar plates containing chloramphenicol 5 µg/mL and +/− aTc 20ng/mL and incubated overnight at 30°C.(B) Overnight cultures grown with aTc were sub-cultured in 1:1000 dilution ratio into fresh 5 mL LB ± aTc and grown at 37°C for 3hrs, then sub-cultured again into LB ± aTc media. The O.D._600_ of the cultures were measured at 3 and 6 hours, and the ratio of Δ6LTG *P_aTc_-dsvR* to WT *P_aTc_-dsvR* in ± aTc conditions was plotted as a bar graph. Statistical significance was determined with Welch’s two-sample t-test. *, p < 0.05; ns, p >= 0.05

**Figure 3:**
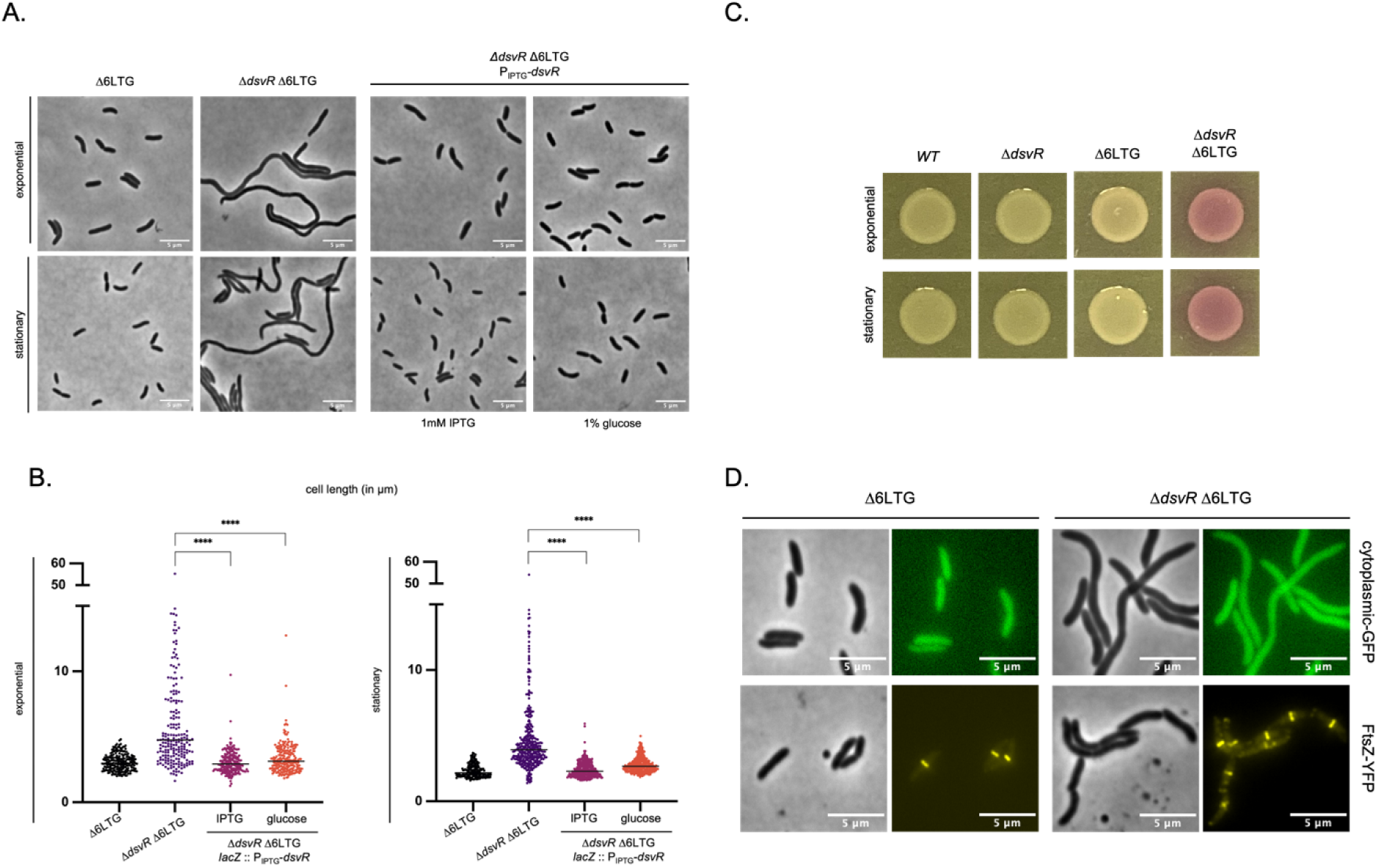
Deletion of *dsvR* in Δ6LTG background leads to severe cell filamentation and lysis. (A) Overnight cultures were 1:100 sub-cultured into LB and grown at 37°C until they reached the exponential/stationary phase and were imaged using phase contrast microscopy. The strain Δ*dsvR* Δ6LTG P_IPTG_-*dsvR* was grown in LB + IPTG/glucose media. Scale bar: 5 µm. (B) Scatter plots showing length distributions of cells. The cells were segmented using Omnipose, analyzed using MicrobeJ in ImageJ, and plots were generated in GraphPad Prism. Statistical significance was determined with Welch’s two-sample t-test. ****, p < 0.0001. (C)To visualize lysis, 10µL overnight cultures were spotted on LB-agar plates containing CPRG 120 µg mL^−1^ and incubated overnight at 30°C. (D) Overnight cultures expressing cytoplasmic GFP or FtsZ-YFP from from an IPTG-inducible promoter were 1:100 sub-cultured in LB containing 1mM IPTG, grown at 37°C until they reached the exponential, and were imaged using phase contrast and fluorescence microscopy. Scale bar: 5 µm. causes dramatic cell division defects upstream of septation in LTG deficient cells.

### The DsvR regulon includes catabolic and cell division functions

DsvR is a homolog of SgrR in *E. coli* to which it shows a 54% similarity. The Alphafold predicted structure of DsvR shows an N-terminal HTH domain, and a C-terminal SBP_bac5 solute-binding domain (Fig. S2). In enteric bacteria like *E. coli* and *Salmonella typhimurium*, SgrR controls expression of itself, the small protein SgrT, and the regulatory small RNA SgrS. SgrS and SgrT subsequently differentially regulate a glucose phosphate stress response regulon. Most notably, this includes production of a sugar phosphate phosphatase YigL, the sugar efflux pump SetA, and post-transcriptional downregulation of the primary glucose permease PtsG ^49–52^. Importantly, *V. cholerae’s* genomic context of *dsvR* is very different from *sgrR,* as *V. cholerae* lacks the small RNA SgrS homolog upstream of the transcription factor, and also the SetA efflux pump ^53,54^. This raises the possibility that DsvR fulfills a different function in *V. cholerae*. We therefore conducted an RNAseq experiment to determine the DsvR regulon.

Since we observed a dramatic phenotype in the Δ6LTG genetic background, indicating that the DsvR regulon was both expressed and important under these conditions, we first compared global transcript abundances between the Δ*dsvR* Δ6LTG mutant and its Δ6LTG parent. The DESeq2 RNA-Seq analysis revealed a list of differentially expressed genes using the cutoffs |log_2_-fold change| > 2.5 and p-value < 0.05 (Fig. 4A). Nutrient acquisition and catabolism functions were notably differentially expressed, including several carbon source uptake and catabolism genes that were markedly upregulated in the mutant (indicating negative regulation by DsvR). This included the RbsACDK ribose ABC transporter and kinase system (*vca0127 – vca0131)*, which provides a precursor for the non-oxidative branch of the pentose phosphate pathway (PPP); trehalose uptake and catabolism (*vc0910 – vc0911*); gluconate permease and gluconate kinase (feeding into the Entner-Doudoroff (ED) pathway and, through the gluconate kinase that was also upregulated in the mutant, pentose phosphate pathway (PPP)); as well as *malF* and maltoporin (*lamB*). In contrast, several genes were putatively positively regulated by DsvR (i.e., reduced expression in the mutant). This included the operon encoding the DNA uptake pilus; an uncharacterized putative sodium/solute symporter system (*vc2704* – *vc2705*); chitodextrinase (*vca0700*) and endo-chitodextinase (*vca0680*) involved in chitin catabolism; and the *mglBAC* (*vc1325* – *vc1328*) uptake system for galactose/glucose. Thus, the majority of differentially regulated genes seem to indicate that DsvR orchestrates a response that favors substrates that feed into the glycolysis pathway (chitin, maltose, galactose/glucose) over those that feed into ED/PPP.

**Figure 4:**
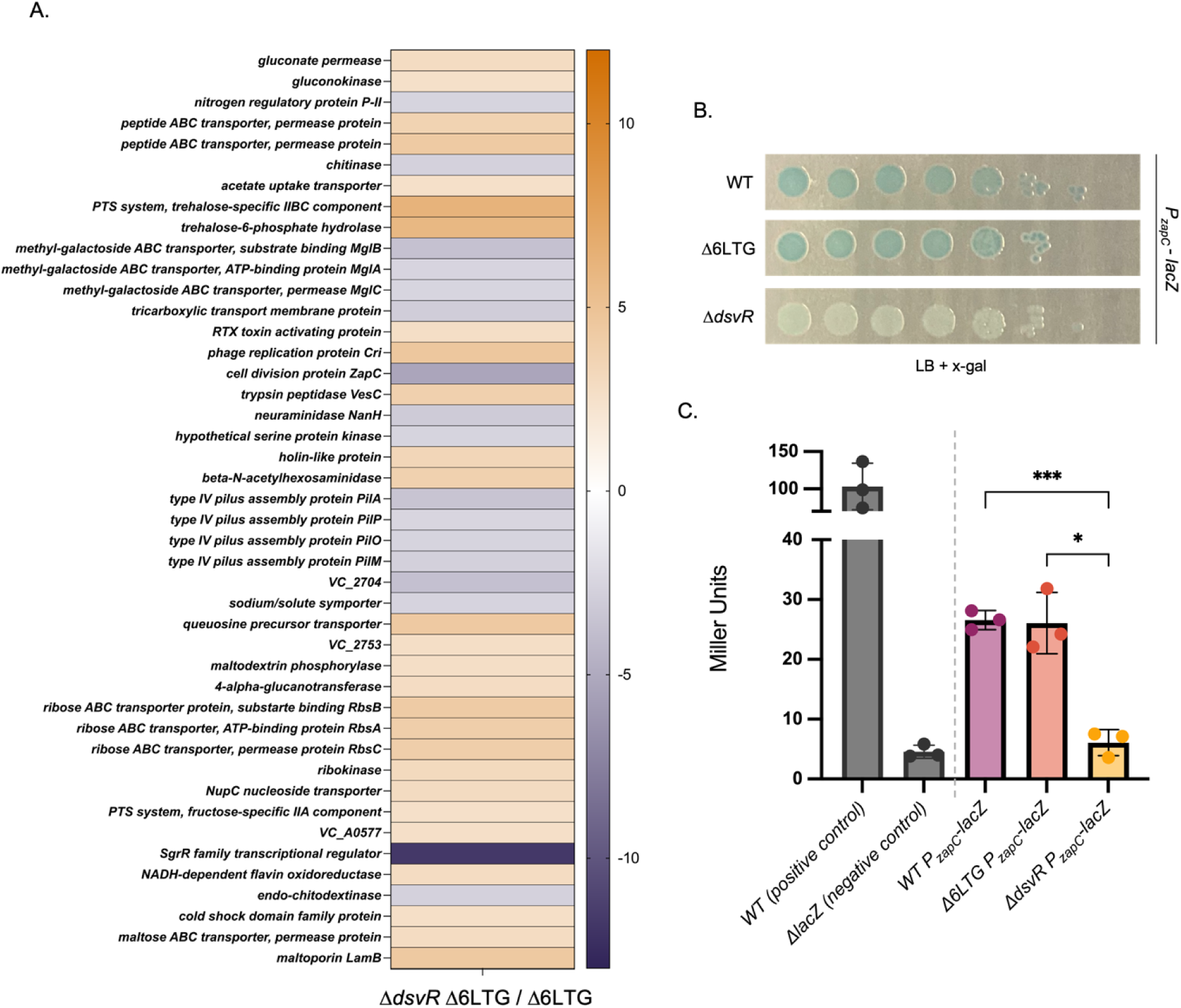
DsvR positively regulates the induction of the gene *zapC*. (A) RNASeq heatmap showing mRNA transcripts differentially expressed (where log_2_fold-change > 2.5 or log_2_fold-change < 2.5, p-adj < 0.05) in Δ*dsvR* Δ6LTG normalized to the Δ6LTG control strain. Purple colour indicates genes positively regulated by DsvR; orange colour indicates genes negatively regulated by DsvR; (B) The overnights of strains containing *lacZ*-transcriptional fusion of *zapC* in WT, Δ6LTG or Δ*dsvR* background were serially diluted from 10^0^ to 10^−7^ in LB, spotted (5µL) on LB-agar plates containing X-gal 120 µg/mL, and incubated overnight at 37°C; (C) A bar-graph depicting the quantification of the *P_zapC_-lacZ* promoter activity using Miller Units by measuring β-galactosidase (LacZ) activity against an ONPG chromogenic substrate. WT, Δ6LTG or Δ*dsvR* strains containing the *P_zapC_-lacZ* construct were grown overnight and diluted 1:100 into LB and grown in 30°C until the O.D._600_ reached 0.5 for LacZ activity quantification. The bar graph was generated in GraphPad Prism. Statistical significance was determined with Welch’s two-sample t-test. ***,p < 0.001; *, p < 0.05

Finally, one gene was not like the others: Unexpectedly, the most strongly upregulated gene in the putative DsvR regulon was the cell division gene *zapC.* It encodes a non-essential cell-division protein that stabilizes the Z-ring by suppressing GTPase activity and promoting lateral interactions of FtsZ polymers *in vitro* ^16,17^. Transcriptional regulation of cell division functions was intriguing, particularly in the context of our subtly division-defective Δ6LTG strain. A connection between *dsvR* and *zapC* was further solidified by our TnSeq dataset, as *zapC,* like *dsvR,* was conditionally essential in Δ6LTG (Fig. 1C).

### DsvR positively regulates the induction of *zapC* transcription

To validate whether DsvR controls *zapC* expression, we constructed a *zapC* transcriptional fusion (P*_zapC_-lacZ*) by fusing its promoter region with the *lacZ* reporter gene, when expressed forms blue coloration when grown on media containing the chromogenic substrate X-gal. In both the WT and Δ6LTG background, we observed equal blue coloration, indicating the same level of constitutive expression in both strains (Fig. 4B, C). This was corroborated by our RNASeq experiment, which revealed lack of differential expression of DsvR regulon members between WT and Δ6 LTG (Fig. S3). However, P*_zapC_* activity was significantly reduced in the Δ*dsvR* background (Fig. 4B), and restored by overexpression of *dsvR* from a plasmid (Fig. S4), confirming that DsvR positively controls *zapC* expression. DsvR-dependent regulation of *zapC* was also quantified using a Miller assay for β-galactosidase activity (Fig. 4C).

### Identification of the putative DsvR binding box

While trying to identify the putative DsvR binding box in the promoter of *zapC*, we noticed an 11bp sequence in P*_zapC_* that was identical to a 11bp sequence present immediately upstream of the *dsvR* ORF start codon, prompting us to propose that this was likely the transcription factor binding box in P*_zapC_* (Fig. 5A). In *E. coli*, SgrR negatively auto-regulates itself ^50^. To test if this sequence affected *zapC* promoter activity, we deleted the 11bp from the P*_zapC_-lacZ* construct (hence designated as *P_zapC(Δ11)_*), and plated strains carrying those constructs on agar containing X-gal. Consistent with an important role in promoter activation, blue coloration was abolished in the WT carrying this *P_zapC(Δ11)-_lacZ* construct, suggesting a decline in P*_zapC_* activity upon deletion of the putative binding site.

**Figure 5:**
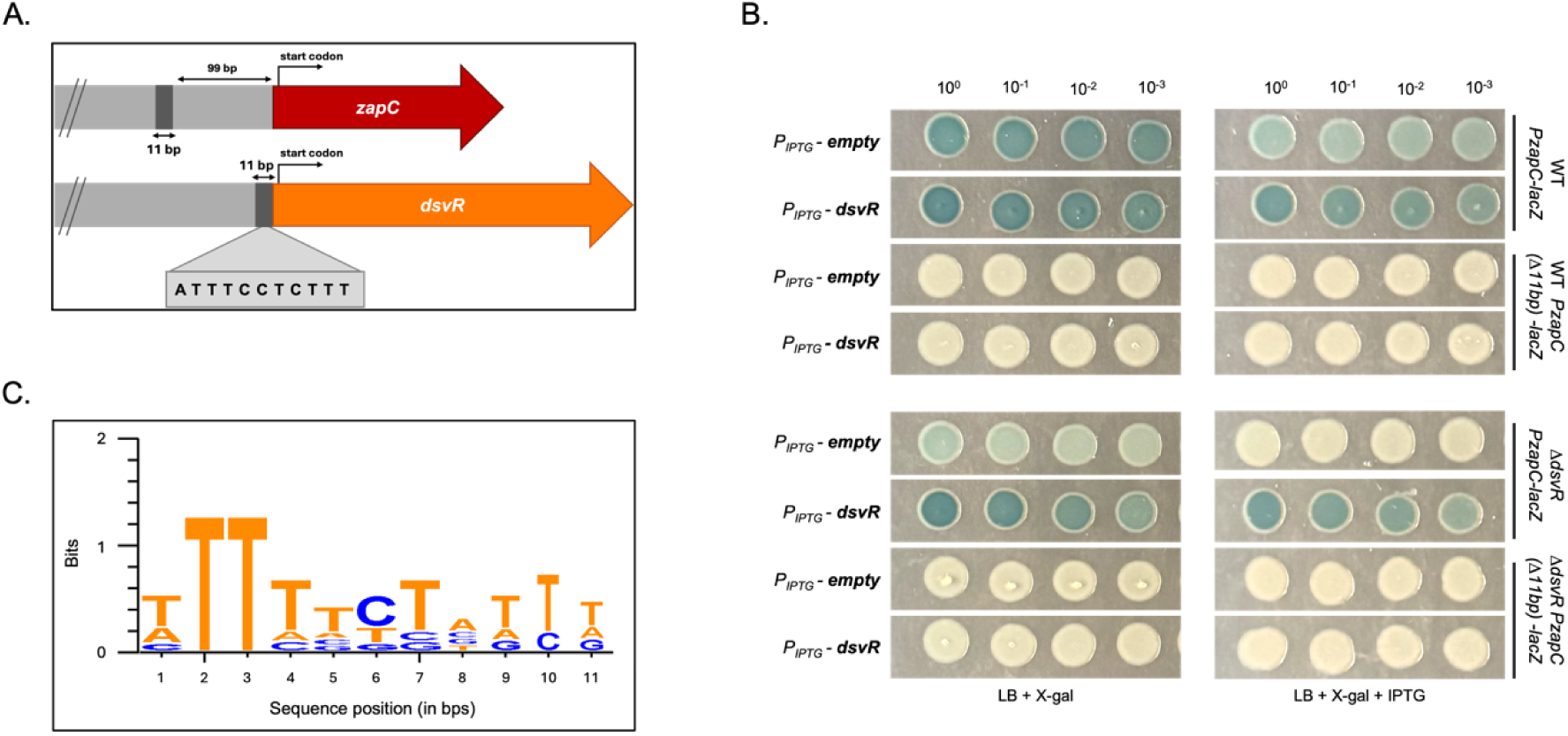
The *zapC* promoter contains a putative transcription factor binding box. (A) Schematic showing an 11bp sequence conserved between the *zapC* and *dsvR* promoters; (B) Strains where 11bp were deleted from the *P_zapC_* region (now designated as *P_zapC(Δ11)_* - *lacZ*) carried either a control pHL100 plasmid or over-expressed *dsvR* from an IPTG-inducible promoter. The overnights were serially diluted from 10^0^ to 10^−3^, spotted (5µL) on LB-agar plates containing X-gal 120 µg/mL, +/− 1mM IPTG, and incubated overnight at 37°C; (C) Sequence logo showing the consensus putative transcription factor binding box found in the promoter of *zapC*. The promoters of 6 genes/operons positively regulated by DsvR (from Fig. 4A) were aligned using MUSCLE in SnapGene, and the sequence logo was generated in WebLogo3.

Similarly, the Δ*dsvR* strain exhibited negligible P*_zapC_* and P*_zapC(Δ11)_* activity. Additionally, in contrast to the P*_zapC_-lacZ* strains, *P_zapC(Δ11)-_lacZ* activity was not restored by overexpression of *dsvR* from an IPTG-inducible promoter using a multicopy plasmid (Fig. 5B). We further aligned the putative promoter of *zapC* with the promoters of 6 other genes positively controlled by DsvR (based on RNASeq results) using MUSCLE in SnapGene and found some consensus as shown in the Sequence logo generated with WebLogo3^55^ (Fig. 5C). We also aligned the putative promoter of *zapC* with 25 other promoters controlled (positively or negatively) by DsvR and found a similar conservation of this 11bp sequence region (Fig. S5). We conclude that DsvR-dependent *zapC* expression relies on this putative binding site.

### *zapC* overexpression is sufficient to suppress Δ*dsvR* defects

Our data suggested that DsvR positively regulates *zapC*; we therefore reasoned that the Δ*dsvR* phenotypes might be caused by insufficient *zapC* levels. If this was the case, *zapC* overexpression should rescue these phenotypes. To test this, we placed *zapC* under control of an IPTG-inducible promoter and integrated this construct into a neutral chromosomal locus in the Δ*dsvR* Δ6LTG strain. The strains were then imaged upon induction of *zapC* expression. This revealed a complete restoration of cell length when *zapC* was overexpressed. Partial restoration was observed even in the absence of inducer, showing that even leaky expression from the promoter was sufficient (Fig. 6A, B). Since the ZapC homolog from *E. coli* showed 52% sequence similarity to N16961 ZapC, we used Alphafold3 to predict the structures and interactions between ZapC and FtsZ from *E. coli* and/or N16961. The iPTM and pTM scores for the interaction between *E. coli* ZapC - N16961 FtsZ were comparably high like the native N16961 ZapC - FtsZ pair (Fig. S6). Therefore, we tested cross-species complementation by overexpressing the *E. coli* homolog in the N16961 Δ*dsvR* Δ6LTG mutant, and as expected, this was sufficient to rescue the filamentation phenotype (Fig. 6C). Thus, filamentation in the Δ*dsvR* Δ6LTG background is caused by insufficient *zapC* expression.

**Figure 6:**
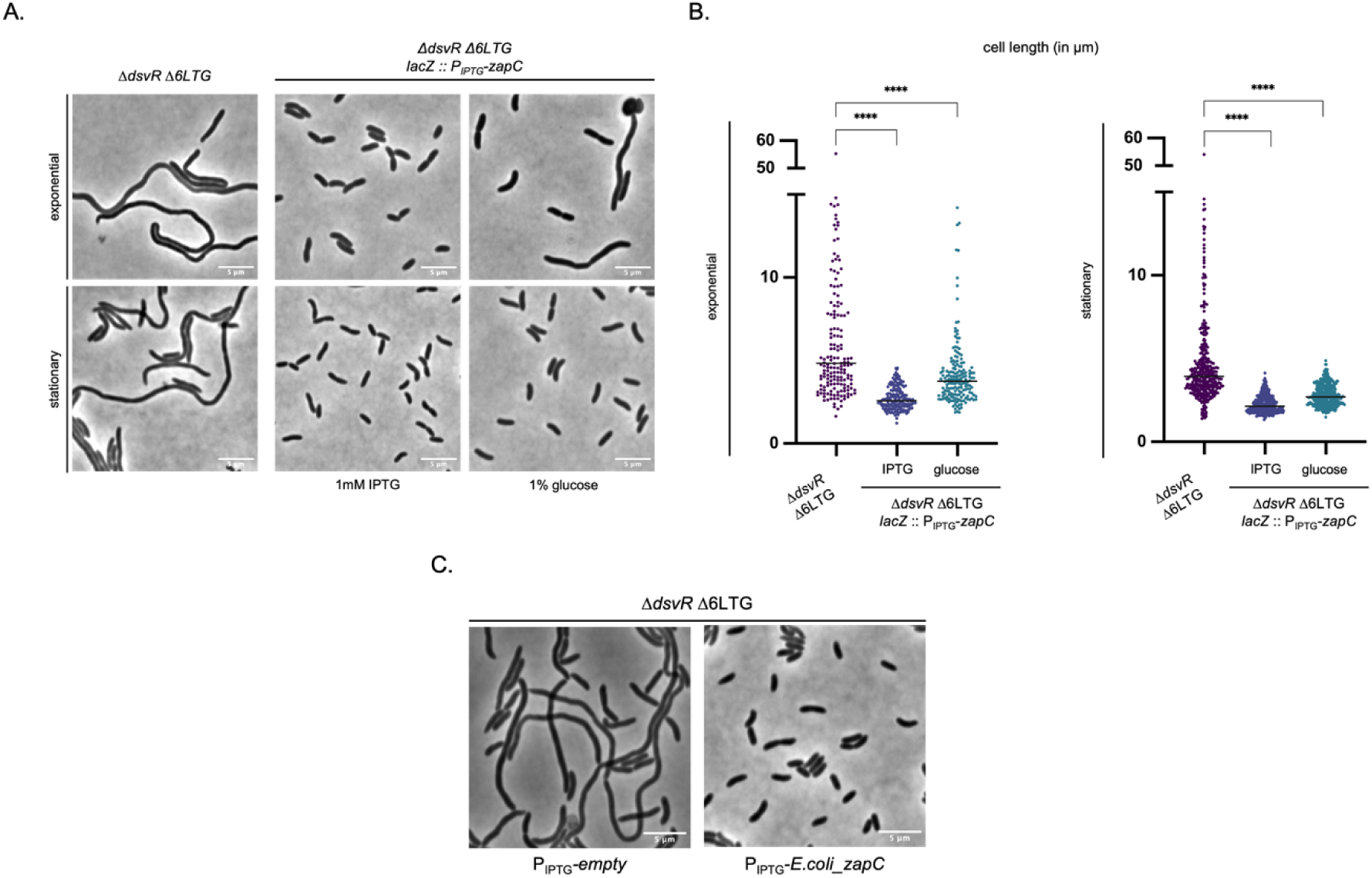
ZapC overexpression rescues cell length in the filamentation mutant. (A) Overnight cultures were 1:100 sub-cultured in LB containing 1mM IPTG or 1% glucose, grown at 37°C they reached the exponential stage and then imaged using phase contrast microscopy. Scale bar: 5µm; (B) Scatter plots showing length distributions of cells. The cells were segmented using Omnipose, analyzed using MicrobeJ plugin in ImageJ, and plots were generated in GraphPad Prism. Statistical significance was determined with Welch’s two-sample t-test. ****, p < 0.0001; (C) Overnight cultures were 1:100 sub-cultured in LB containing 1mM IPTG, grown at 37°C for 2hrs and then imaged using phase contrast microscopy. Scale bar: 5µm.

To dissect this relationship further, we attempted to delete the *zapC* gene in the Δ6LTG background. Despite multiple attempts, we were unable to do so (suggesting true essentiality, as also indicated by our TnSeq) and opted for a depletion approach instead. To this end, we replaced the native *zapC* promoter with an anhydro-tetracycline (aTc)-inducible promoter in both WT and Δ6LTG. As expected, *zapC* depletion in the Δ6LTG background (but not WT) resulted in a dramatic filamentation and lysis (Fig. S7B), reduced growth in liquid medium, and a pronounced plating defect (Fig. 7A, B). All these phenotypes were reminiscent of *dsvR* depletion, albeit more severe (Fig. S7A).

**Figure 7:**
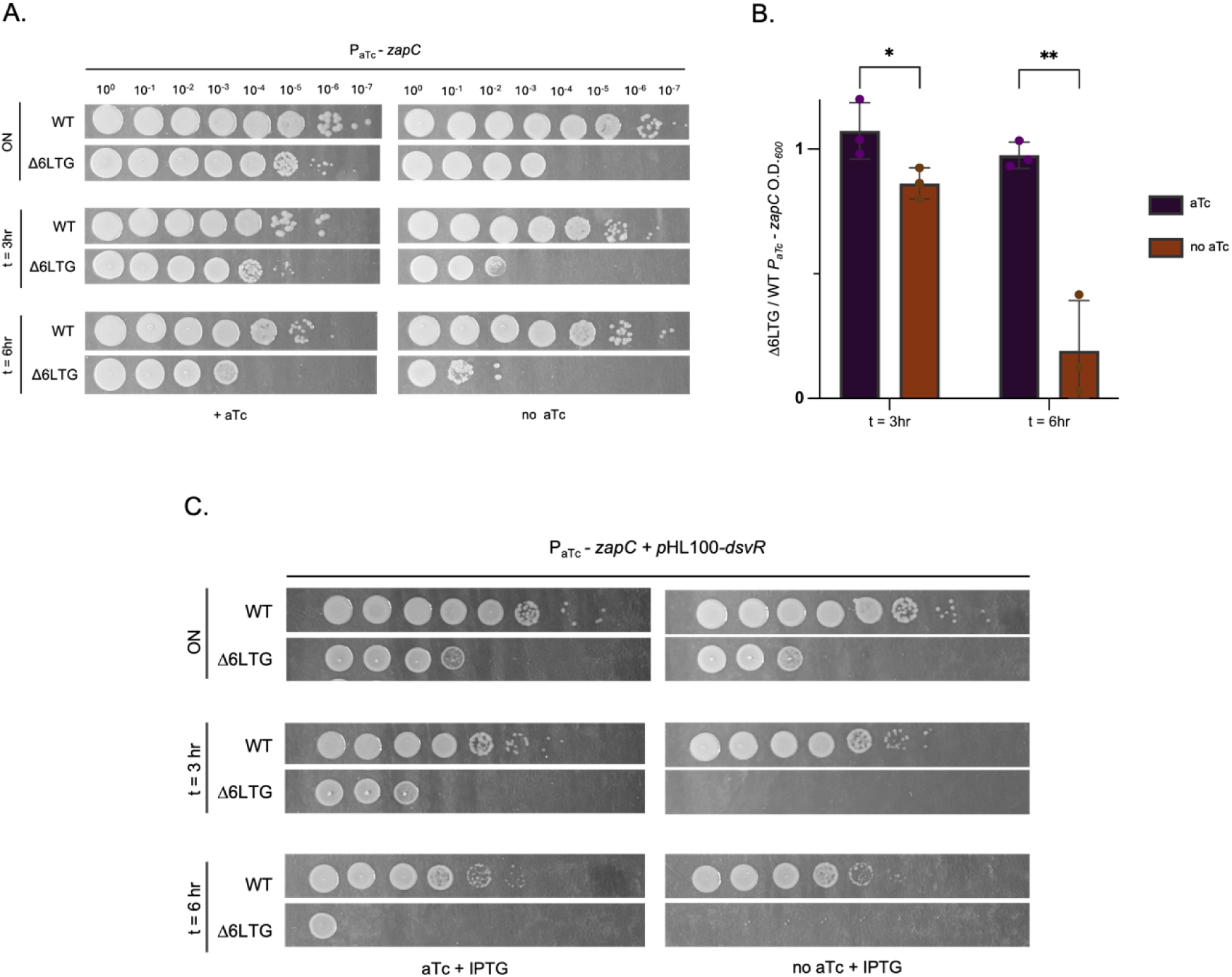
*zapC* acts downstream of *dsvR*. (A) The gene *zapC* was depleted by replacing its native promoter with an anhydrotetracycline(aTc)-inducible promoter. Overnights were sub-cultured in a 1:1000 into LB ± aTc 20 ng/mL and grown at 37°C for 3hrs, then sub-cultured 1:1000 and grown for another 3hrs. At mentioned time-points, they were serially diluted, and 5µL was spotted on LB-agar plates containing chloramphenicol 5 µg/mL and +/− aTc 20ng/mL and incubated overnight at 30°C; (B) Overnights were sub-cultured as described above. O.D._600_ was measured at 3 and 6 hours, and the ratio of Δ6LTG *P_aTc_-zapC* to WT *P_aTc_-zapC* was plotted as a bar graph in GraphPad. Statistical significance was determined using Welch’s two-sample t-test. *, p < 0.05; **, p < 0.01; (C) Overnights were sub-cultured in a 1:100 dilution ratio into LB ± aTc 20ng/mL ± 1mM IPTG, and grown at 37°C for 3hrs, then sub-cultured again in 1:100, and grown for another 3hrs. Cultures were serially diluted at indicated time-points and 5µL was spotted on LB-agar plates containing chloramphenicol 5 µg/mL, kanamycin 50 µg/mL, +/− aTc 20ng/mL, +/− 1mM IPTG, and incubated overnight at 30°C.

We next conducted more detailed epistatic analysis. If *zapC* operates in the same pathway as, and downstream of, *dsvR,* then not only should overexpressing *zapC* restore Δ*dsvR* Δ6LTG phenotypes (shown in Fig. 6), but overexpression of DsvR should also fail to rescue the absence of *zapC*. To test this, *dsvR* was expressed ectopically from a plasmid with IPTG while depleting *zapC* using the aTc system described above. When *zapC* was depleted, Δ6LTG cells failed to grow despite *dsvR* overexpression (Fig. 7C, S8). Collectively, these results suggest that *dsvR* and *zapC* act in the same pathway, where *zapC* expression is primarily controlled by *dsvR*.

### FtsZ abundance is limiting for cell division in LTG-deficient strains

Our data thus far indicated that DsvR-induced *zapC* is essential for proper cell division upon LTG deficiency. We hypothesized that this was due to ZapC’s absence affecting Z-ring stability. If that is the case, increasing FtsZ abundance should rescue phenotypes caused by absence of *dsvR*. To test this, we turned to the cell division defect in a Δ*dsvR* Δ6LTG mutant. Overexpression of *ftsZ* from an IPTG-inducible promoter was sufficient to restore cell length (Fig. 8A). While testing the effect of overexpression of other divisome components, we observed that the Z-ring stabilizers ZipA and FtsA had an adverse but expected phenotype of further inhibition of cell division, thus aggravating the filamentation. On the other hand, FtsN overexpression also rescued the Δ*dsvR* Δ6LTG filamentation mutant (Fig. S12). Overall, these data suggest that the abundance of the structural cytoskeletal homolog FtsZ and the late divisome protein FtsN can obviate the need for ZapC’s Z-ring stabilizing function in the Δ*dsvR* Δ6LTG mutant.

**Figure 8:**
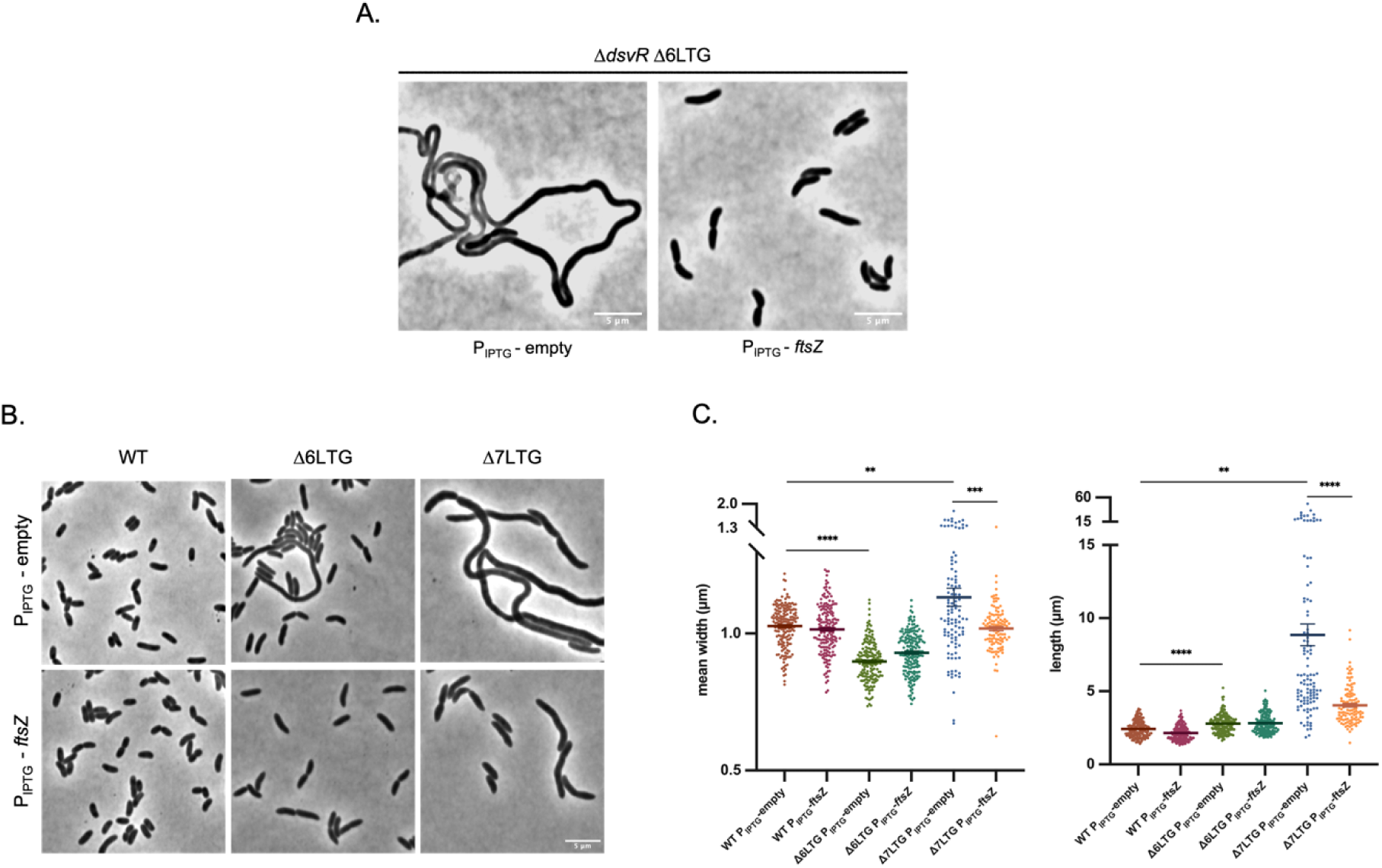
FtsZ overexpression improves length-width homeostasis of rapidly growing LTG deficient mutants. (A) Overnight cultures were 1:10,000 sub-cultured in LB containing 1mM IPTG and grown at 37°C they reached the exponential stage (about 3.5hrs) and then imaged using phase contrast microscopy. The strains WT, Δ6LTG or Δ7LTG contained either P_IPTG_-empty or P_IPTG_-*ftsZ* constructs. Scale bar: 5µm. (B) Scatter plots showing mean width and length distributions of exponentially growing cells. The cells were segmented using Omnipose and analyzed using MicrobeJ plugin in ImageJ. Statistical significance was determined with Welch’s two-sample t-test. **, p < 0.01; ***,p < 0.001; ****, p < 0.0001; ns, p >= 0.05

To investigate the role of FtsZ abundance in LTG definiciency phenotypes further, we used an even more minimal LTG strain (Δ7LTG), where RlpA is the only remaining LTG, and assessed the role of *zapC* in proper growth and morphogenesis in this background. We previously found (and corroborated here) that LTG-deficient mutants, particularly Δ7LTG, exhibit severe morphology defects (increased length, width, and viability defects) in a dilution-dependent manner (Fig. 8B) ^40^. These defects were most pronounced when cultures spent prolonged time in exponential phase, likely caused by growth-dependent periplasmic accumulation of PG breakdown products ^40^. We hypothesize that this makes division more challenging, as Z-ring force will have to overcome periplasmic debris, which perhaps causes perturbations in PG synthesis. To test if FtsZ limitation might contribute to these phenotypes, we overexpressed *ftsZ* in the Δ7LTG background, and measured growth and morphology during extended exponential phase (achieved via back-dilution). Interestingly, FtsZ overexpression vastly improved the morphology of the Δ7LTG (Fig. 8B, C, S9). These observations suggest that FtsZ abundance and/or stability is limiting for cell division in ΔLTG strains.

### Δ*dsvR* and Δ*zapC* mutants are sensitive to rod-complex inhibition

Our data so far support a model where ZapC fortifies Z-ring function under adverse conditions like LTG insufficiency. We reasoned that the DsvR-ZapC pathway might more broadly promote cell division when cell size and shape are disturbed. To test this, we interrogated the role of this pathway in intrinsic resistance to cell wall-acting antibiotics. Plating experiments revealed that Δ*dsvR* and Δ*zapC* were hyper-sensitive to inhibitors of the rod-complex, specifically A22 and mecillinam (Fig. 9A), but, perhaps counter-intuitively given ZapC’s demonstrated role in division, not sensitive to aztreonam (inhibits divisome component FtsI), nor to moenomycin (inhibits aPBPs) (Fig. S10 A, B). A22 inhibits polymerization of MreB^56^, the cytoskeletal component that associates with the rod-complex to promote PG insertion, whereas mecillinam is a β-lactam antibiotic that inhibits the transpeptidase activity of PBP2 ^57^. Sensitivity on plates was recapitulated in liquid LB medium, where the Δ*dsvR* and Δ*zapC* mutants exhibited a slight decrease in optical density in the presence of either mecillinam or A22 (Fig. 9B). This could be indicating that Δ*dsvR* and Δ*zapC* mutants undergo fewer cell divisions than WT. Conversely, overexpression of this pathway promoted growth and increased resistance to these antibiotics in the WT background: *zapC* overexpression improved survival in both solid and liquid media, whereas *dsvR* overexpression improved growth only in liquid media (Fig. 9A, B).

**Figure 9:**
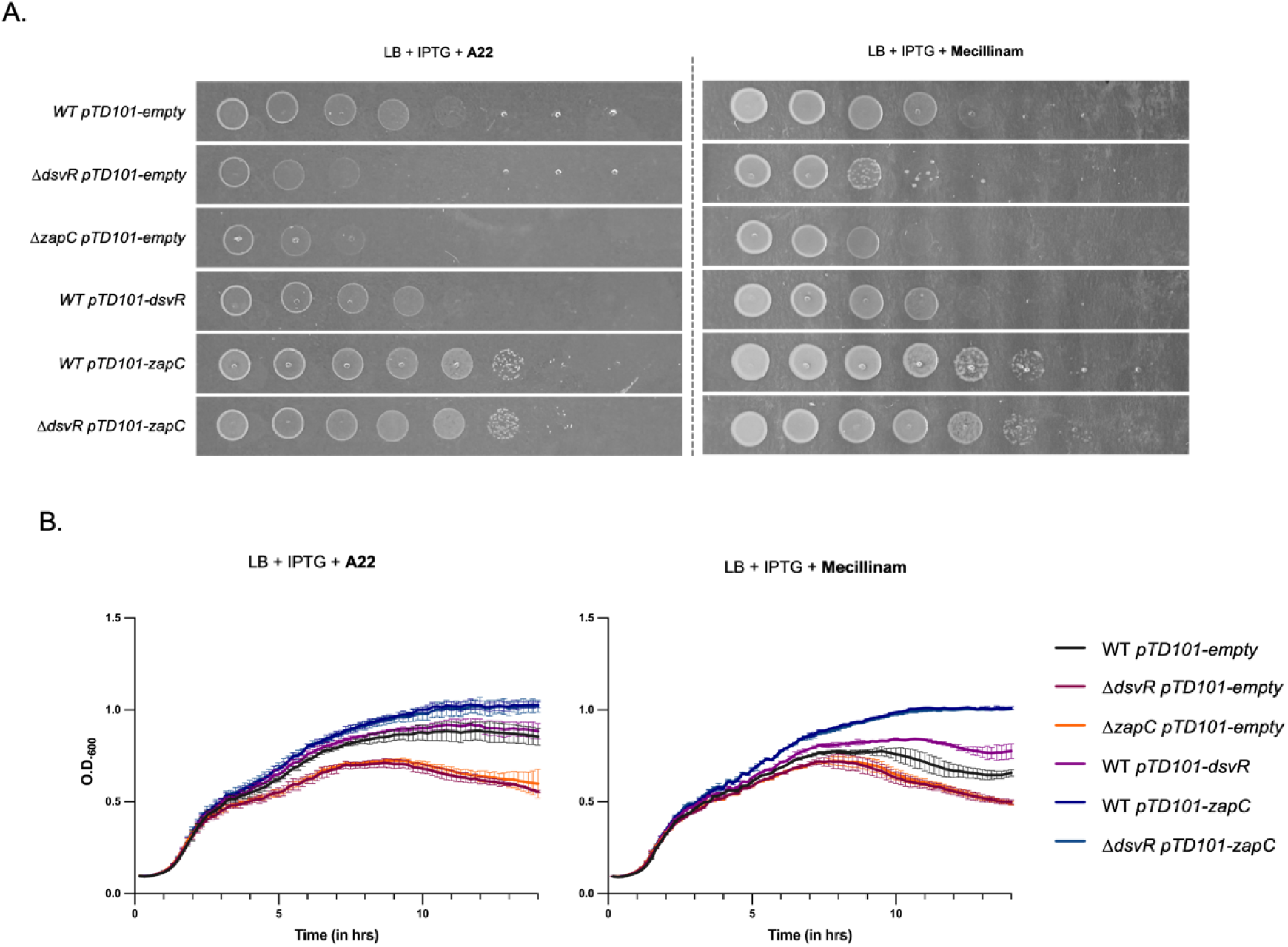
Δ*dsvR* and Δ*zapC* mutants are sensitive to rod-complex inhibition, and overexpression of these can render resistance. (A) Representative growth curves of strains grown in LB containing 1mM IPTG, and A22 10 µg/mL or Mecillinam 2 µg/mL. Overnights were diluted 1:100 into LB media and their O.D._600_ was recorded over time, shaking at 37°C in a honeycomb plate in a Bioscreener machine; (B) Saturated overnight cultures harboring chromosomal IPTG-inducible overexpression constructs were serially diluted from 10^0^ to 10^−7^ in LB, and 5µL was spotted on LB-agar plates containing 1mM IPTG, A22 2.5 µg/mL or Mecillinam 0.5 µg/mL and incubated overnight at 30°C.

Given the prominent role of ZapC in cell division, we were surprised by the enhanced sensitivity to rod-complex inhibitors, rather than those affecting division. However, a well-characterized mechanism of resistance against rod-complex inhibition relies on FtsZ overexpression ^58^; this is likely due to cell width increasing to a point where the Z-ring cannot form efficiently, which can be overcome by increasing FtsZ abundance. We thus hypothesized that ZapC stabilizes the Z-ring to promote its efficiency of closing under increased cell width conditions. If this is the case, WT cells should be able to undergo more divisions than Δ*zapC* in the presence of these rod-complex inhibiting antibiotics. To test this, we grew WT and Δ*zapC* strains in the presence of A22 or mecillinam. After 30 minutes of exposure, inhibition of rod-complex induced spherical cells, as expected, but we noted a clear difference in cell size and morphology between Δ*zapC* mutant and WT. The Δ*zapC* cells were more elongated and wider than WT (Fig. 10 A, B), indeed suggesting that during rod-complex inhibition, division stops earlier in Δ*zapC*. We next conducted single-cell quantification of growth dynamics on agarose pads containing A22. Time-lapse experiments of WT vs. Δ*zapC* and further analyses revealed that the Δ*zapC* mutant underwent fewer rounds of division before terminally enlarging (Fig. 10 C, D, E). Thus, ZapC promotes cell division under otherwise unfavorable conditions, likely through stabilizing the Z-ring. Interestingly, ZapC/DsvR appear to play a role in length as well as width homeostasis even in the absence of antibiotic stress, as we found that the Δ*dsvR* and Δ*zapC* mutants were both longer and wider than WT (Fig. S11 A,B). This perhaps points to an interplay between regulation of cell division and elongation, as suggested previously. ^59,60^

**Figure 10:**
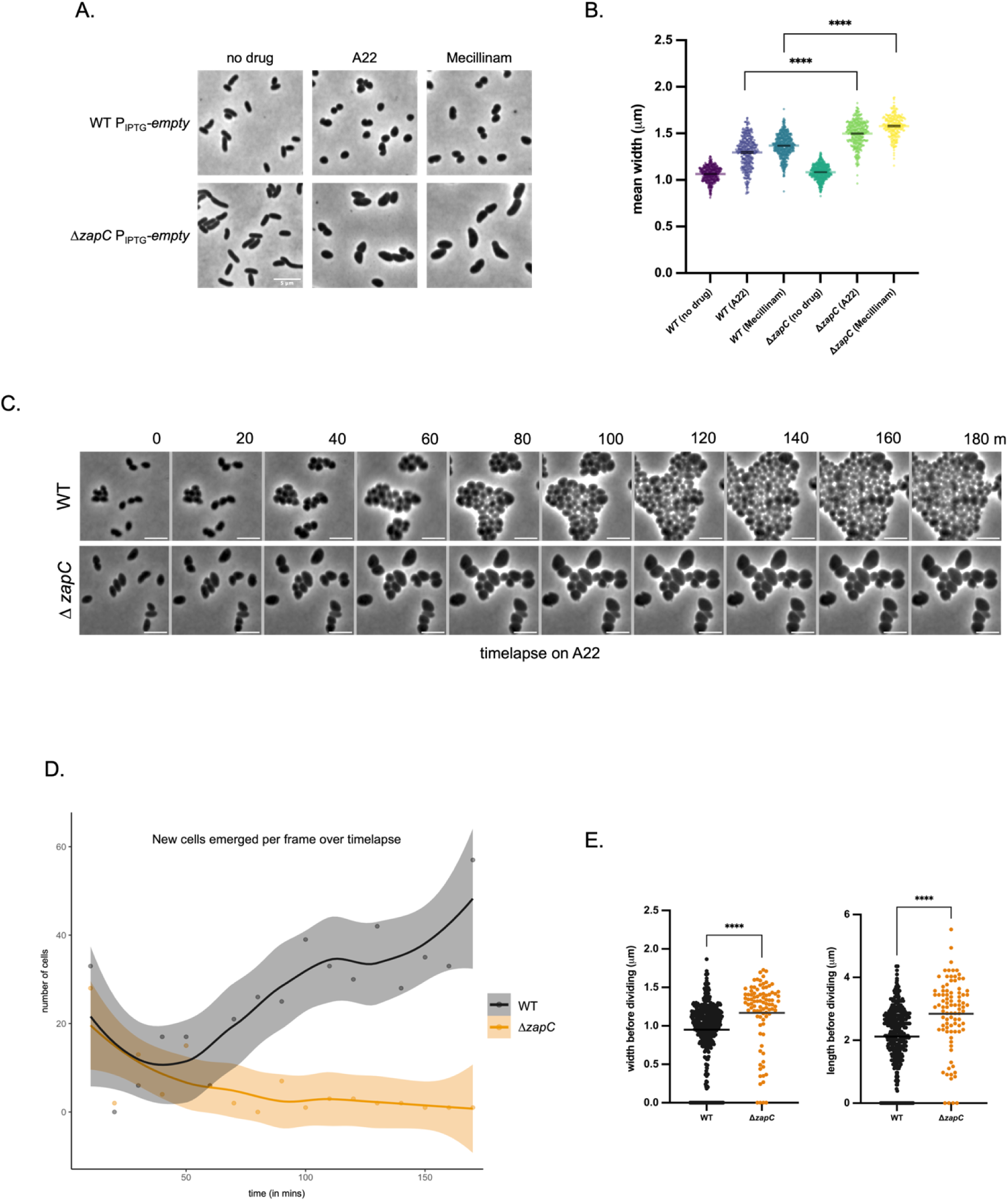
Δ*zapC* undergoes fewer rounds of cell division and is larger than WT during rod-complex inhibition. (A) Overnight cultures were 1:100 sub-cultured in LB containing 1mM IPTG and grown at 37°C for 1.5 hours. To these, A22 20 µg/mL or Mecillinam 5 µg/mL was added, and these were further grown for another 30mins, then imaged using phase contrast microscopy. Scale bar: 5µm; (B) Scatter plots showing mean width distributions of strains grown in antibiotics as mentioned above. The cells were segmented using Omnipose and analyzed using MicrobeJ plugin in ImageJ, plots were generated in GraphPad Prism. Statistical significance was determined with Welch’s two-sample t-test. ***,p < 0.001; ****, p < 0.0001; (C) A montage showing phase contrast time-lapse of cells grown on LB+0.8% agarose pads containing A22 10 µg/mL. Scale bar: 5µm; (D) Line plots showing number of new cells generated in each frame over time during the time-lapse. The movies were analyzed using OmniSegger in MATLAB, and the plot was generated in R; (E) Scatter plots showing mean width and length (before a cell disappears from the frame) distributions of all cells over the entire time-lapse. The movies were analyzed using OmniSegger in MATLAB, and plots were generated in GraphPad Prism. Statistical significance was determined with Welch’s two-sample t-test. ****, p < 0.0001

We next hypothesized that similar to the ΔLTG backgrounds, Δ*zapC* sensitivity to rod-complex inhibitors was due to constraints in FtsZ function. To test this, we overexpressed *ftsZ* in WT vs. Δ*zapC* vs. Δ*dsvR* backgrounds. Increasing FtsZ abundance conferred resistance to A22 and mecillinam not only in the WT but also in the Δ*zapC* and Δ*dsvR* mutants (Fig. 11 A, B). The WT results partially recapitulate the data obtained in *E. coli*, where FtsQAZ overexpression confers A22 resistance, and conditional mecillinam resistance. ^58^ Taken together, our data demonstrate that either ZapC or FtsZ abundance can modulate the ability to divide at increased cell width.

**Figure 11:**
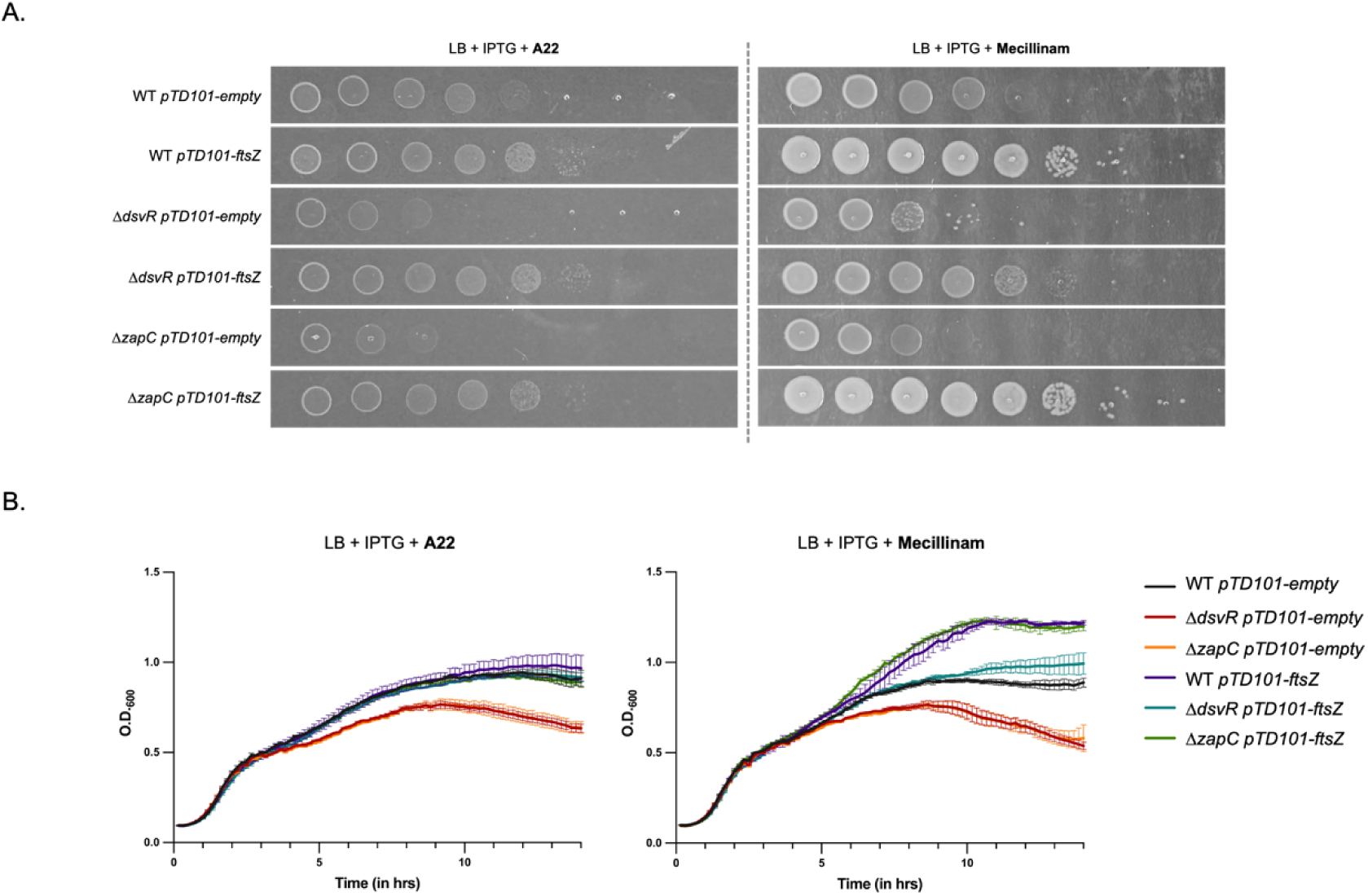
FtsZ overexpression rescues rod-complex sensitivity of Δ*dsvR* and Δ*zapC* and promotes division in the filamentation mutant. (A) Representative growth curves of strains grown in LB containing 1mM IPTG, and A22 10 µg/mL or Mecillinam 2 µg/mL. Overnights were diluted 1:100 into LB media and their O.D._600_ was recorded over time, shaking at 37°C in a honeycomb plate in a Bioscreener machine; (B) Saturated overnight cultures harboring chromosomal IPTG-inducible overexpression constructs were serially diluted from 10^0^ to 10^−7^ in LB, and 5µL was spotted on LB-agar plates containing 1mM IPTG, A22 2.5 µg/mL or Mecillinam 0.5 µg/mL and incubated overnight at 30°C; (C) Overnight cultures were 1:100 sub-cultured in LB containing 1mM IPTG and grown at 37°C for 2 hours and then imaged using phase contrast microscopy. Scale bar: 5µm.

### The absence of aPBP activity partially rescues rod-complex inhibitor sensitivity of *Δ*zapC

The aPBPs likely continue sidewall synthesis during rod-complex inhibition and should thus be limiting for expansion under these conditions. We reasoned that if ZapC’s role is indeed to maintain optimum cell size under which native FtsZ levels accomplish division, the *zapC* mutant’s hypersensitivity to rod-complex inhibition should be suppressed by diminishing aPBP activity, essentially slowing down expansion so that the Z-ring can “catch up” to complete division even during rod-complex inhibition. To test this idea, we deleted *V. cholerae’*s principal aPBP *pbp1A*, or its paralog *pbp1B,* in the Δ*zapC* background, and measured antibiotic sensitivity. Deletion of *pbp1a* improved viability of Δ*zapC* upon mecillinam treatment, increasing plating efficiency >10,000-fold (Fig. 12A). In contrast, deleting the minor aPBP, *pbp1b*, did not have an effect. Next, we used the antibiotic moenomycin that inhibits glycosyltransferase activity of aPBPs. The Δ*dsvR* and Δ*zapC* mutants grew better on LB agar plates that contained sub-MIC moenomycin in addition to mecillinam (Fig. 12B). The overall trend remained conserved upon A22 treatment, albeit the rescue was less robust than using mecillinam (Fig. S13). Microscopic analysis of these strains revealed that co-treatment of mecillinam with moenomycin reverts cell morphology of Δ*zapC* cells back to WT-like (Fig. S14). Collectively, our data suggest that reduction of aPBP activity in Δ*zapC* can promote cell division and growth upon rod-complex inhibition.

**Figure 12:**
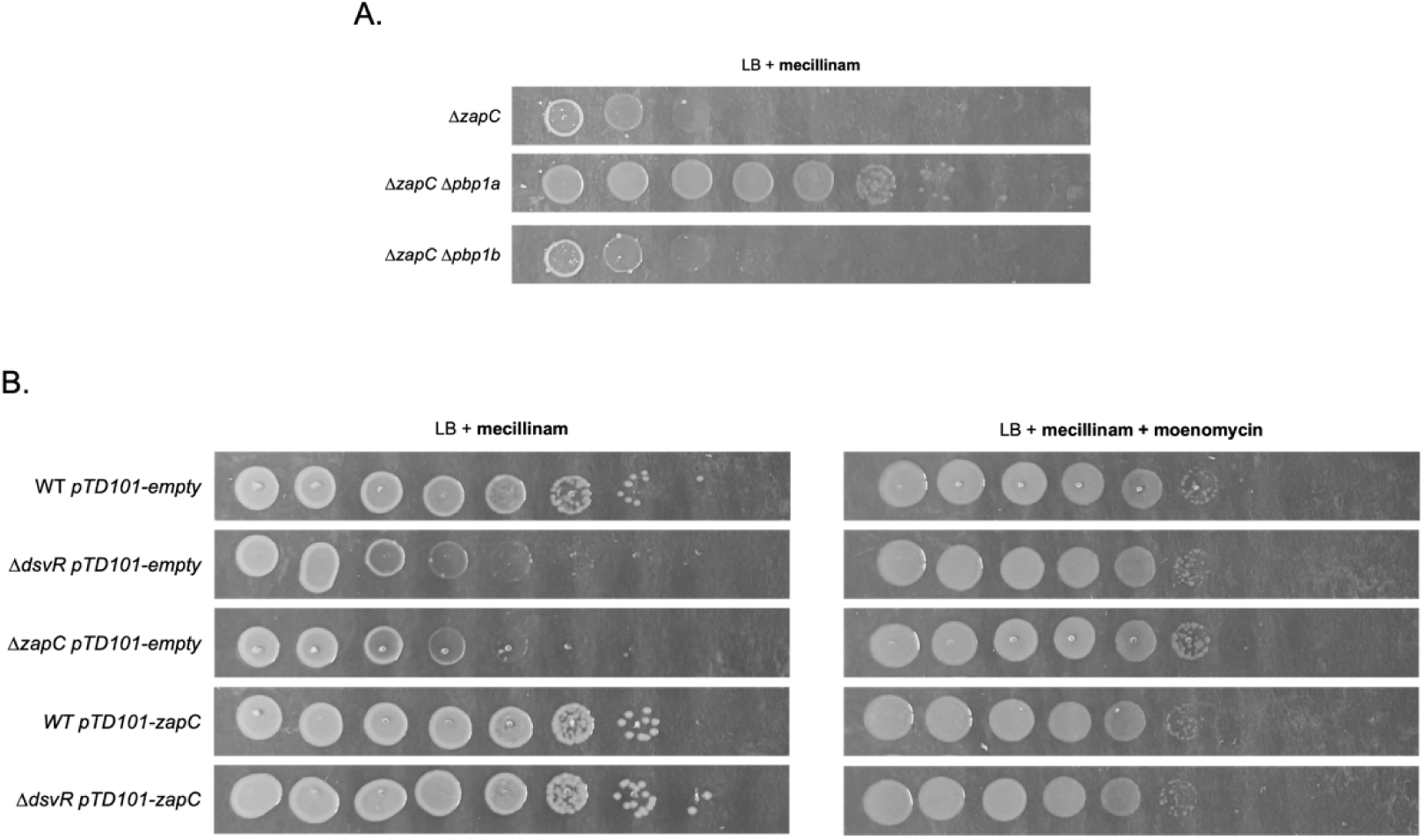
Reduction in aPBP activity rescues mecillinam sensitivity of Δ*dsvR* and Δ*zapC* mutants. (A) Saturated overnight cultures were serially diluted from 10^0^ to 10^−7^ in LB, and 5µL was spotted on LB-agar plates containing mecillinam 1 µg/mL and incubated overnight at 30°C; (B) Saturated overnight cultures harboring chromosomal IPTG-inducible overexpression constructs were serially diluted from 10^0^ to 10^−7^ in LB, and 5µL was spotted on LB-agar plates containing 1mM IPTG, mecillinam 0.5 µg/mL and/or moenomycin 0.5 µg/mL and incubated overnight at 30°C.

## DISCUSSION

Despite intensive work in the field, the roles and regulation of LTGs remain elusive primarily due to two reasons: functional redundancy under laboratory conditions on the one hand, and collective essentiality of LTG enzymatic activity on the other. To dissect the contributions of LTGs in morphogenesis, we generated minimal LTG strains, and used the Δ6LTG strain to interrogate processes vital for growth during LTG insufficiency. In this study, we have shown the importance of cell division and Z-ring stability in ΔLTG strains, and identified a regulatory circuit that maintains it. We show that DsvR, a putative transcription factor, positively controls transcription of ZapC, a Z-ring regulatory protein which further promotes cell division. Our work highlights the importance of Z-ring stability especially when cell width increases beyond a certain threshold, and the underappreciated role of ZapC in supporting cell division under these conditions.

Based on our findings, we propose a model where Z-ring constriction is a limiting factor during LTG insufficiency in *V. cholerae*, and one of the ways to regulate Z-ring condensation and stability is via the DsvR-ZapC pathway. This pathway is dispensable in WT but becomes non-trivial during LTG insufficiency, though our data show that DsvR is not specifically activated in the Δ6LTG background. Instead, deleting the gene *dsvR* reduces overall ZapC transcript abundance, thereby leading to severe cell filamentation and lysis in Δ6LTG, whereas deletion of *dsvR* or *zapC* in WT causes only a slight increase in cell length. Increasing the downstream effector concentration, by overexpressing FtsZ, was sufficient to bypass ZapC essentiality during LTG insufficiency in the Δ*dsvR* Δ6LTG double mutant, and during rod system perturbation. ZapC levels thus become critical when cell division becomes more challenging during stress conditions like LTG insufficiency or cell size changes.

### Similar but different: *E. coli* SgrR vs. DsvR

Interestingly, DsvR is annotated as a homolog of the transcription factor SgrR (“**S**u**g**a**r** transport-related **R**egulator”). In *E.coli*, SgrR regulates the transcription of the sRNA *sgrS* and proteins like SgrT and SetA, which together orchestrate a response to glucose-phosphate stress via downregulating glucose import, and promoting efflux of phosphorylated sugars or their hydrolysis to their non-phosphorylated form ^51,52^. However, *V. cholerae* DsvR appears distinct from *E. coli* SgrR in several crucial ways. The genomic architecture of the locus is very different, as *V. cholerae* lacks both SgrS and SetA ^54^. The DsvR regulon is also different from SgrR, as it positively controls uptake of sugars that enter glycolysis (the opposite of SgrR), but also includes other non-sugar-related functions like Pili, and ZapC. As a notable caveat, our RNAseq design did not allow us to assess differentially regulated small RNAs. While SgrS appears to be absent from *V. cholerae*, we cannot exclude that an analogous regulatory system exists. If this is the case, our transcriptomics dataset might miss some indirectly-regulated functions (e.g., small RNA-mediated translational interference without affecting transcript levels), as is the case for SgrS-mediated regulation in *E. coli;* possibly, the DsvR regulon is thus larger than implied here. However, adding to the suspicion that *V. cholerae* does not have a true SgrS-analogous system, we previously showed that a *V. cholerae pgi* mutant, unlike an equivalent *E. coli* mutant, suffers from significant glucose phosphate toxicity, suggesting that *V. cholerae* is intrinsically “diabetic”, and unable to efficiently cope with sugar phosphate stress ^61^. It is thus possible that the DsvR circuit is a basal carbon source homeostasis sensor, and was co-opted by *E. coli* for sugar phosphate stress by usurping it via a small RNA. Indeed, *E. coli* SgrR’s ability to bind glutamate suggests a role beyond sugar phosphate stress sensing in the *E. coli* homolog as well. ^62^ While we do not currently know what the inducing signal for DsvR is, it is likely that it will be a sugar or sugar-like molecule (albeit probably not glucose-phosphate).

### The importance of ZapC in maintaining normal width during cell division

Our work also sheds light on the function of the poorly-characterized division protein ZapC. ZapC is one of several “**Z**-ring **a**ssociated **p**roteins” that in diverse ways interact with and either stabilize or destabilize the Z-ring, likely often in response to specific cellular needs. ZapC is a widely-conserved accessory protein that stabilizes the Z-ring by binding to the FtsZ globular core ^63^. In *E. coli,* ZapC localizes to the mid-cell, suppresses FtsZ GTPase activity, and promotes lateral interactions of FtsZ polymers *in vitro*. Overexpression of *zapC* in *E. coli* leads to abberrant FtsZ locatizion and lethal filamentation ^64^; in contrast, *V. cholerae* can withstand deletion or overexpression of *zapC*. The apparent essentiality of ZapC only during LTG insufficiency and rod-system inhibition highlights a niche requirement of this protein in reinforcing Z-ring function, potentially when the length-width homeostasis is disrupted, and a stable Z-ring is critical for completing division. Our single cell timelapse data of Δ*zapC* upon A22 exposure are consistent with this idea, as Δ*zapC* keeps increasing in size and is able to undergo fewer rounds of division than WT when elongasome function is compromised. Here we propose that Δ*zapC* grows larger because it has insufficient Z-ring stability and cannot divide. One of the ways to curb enlargement of cells was to decrease the aPBP activity by co-treating the cells with A22/mecillinam and moenomycin. Indeed the co-treatment led to better survival and division of the Δ*zapC* mutant on plate, but we did not see a decrease in cell size. As expected, ZapC and FtsZ overexpression rescues rod-complex inhibition, and promotes cell division probably by strengthening the Z-ring. This is consistent with data in *E. coli,* where FtsZ overexpression can suppress the lethal phenotype of the Rod^−^mutants (inactivation of essential components of rod-complex) or during A22 treatment, where the cells keep dividing as mini spheres. ^58^

### Potential connection between metabolism and cell division

It is intriguing that ZapC is under control of a putative metabolic sensor. Several connections between central metabolism and cell division and growth have emerged in recent years. For example, ZapE, another FtsZ-interacting protein interacts with late-stage cell division protein FtsN and succinate dehydrogenase subunit SdhC of succinate dehydrogenase complex, thereby linking tricarboxylic acid (TCA) cycle and electron transport chain with cell division in Uropathogenic *E. coli* (UPEC) ^65^. In addition, both *E. coli* and *Bacillus subtilis* link cell division and nutrient state through other mechanisms (inhibition of Z-ring formation via UDP-Glucose-responsive UgtP in B. subtilis, and OpgH in *E. coli*) ^66,67^, though the regulatory details differ from the DsvR-ZapC circuit described here.

## METHODS AND MATERIALS

### Bacterial strains, media, and growth conditions

All *V. cholerae* strains used in this study are derivates of El Tor N16961 ^68^, and *E. coli* strains are derivates of *E. coli* K-12 MG1655 ^69^. Strains were grown by shaking at 200rpm at 30 or 37°C in Miller Luria-Bertani (LB) broth (tryptone 10g/L, yeast extract 5g/L, sodium chloride 10g/L) or salt-free LB broth (without sodium chloride). 200μg/mL streptomycin was added to media to grow *V. cholerae* (they are streptomycin-resistant). When appropriate, the following antibiotics were used: carbenicillin (100μg/mL), kanamycin (50μg/mL), chloramphenicol (20μg/mL for *E. coli*, 5μg/mL for *V. cholerae*). For overexpression of genes via IPTG-inducible promoter, 1mM IPTG was used for induction, and 1% glucose was added to prevent expression from the promoter where applicable. For envelope integrity assays, LB was supplemented with chlorophenol red-β-D-galactopyranoside (CPRG, 120μg/mL). Transcriptional LacZ reporters activity was visualized on LB supplemented with 5-bromo-4-chloro-3-indolyl-β-D-galactopyranoside (X-gal 120μg/mL) or quantified using β-galactosidase assays (described below).

### Cloning: strain and plasmid construction

A summary of all strains, plasmids and primers used in this study can be found in supplemental file 3. *E.coli* DH5α λ pir was used for general cloning, while *E.coli* SM10 or MFD λ pir were used for conjugation into *V. cholerae* ^70,71^. Plasmids were built using Gibson assembly ^72^ and verified using Sanger or Oxford Nanopore Technology sequencing. The plasmid harboring the *P_zapC(Δ11)_-lacZ* construct was generated by site-directed mutagenesis of the parent pAW61 *P_zapC_-lacZ* plasmid.

Chromosomal in-frame deletions were generated using the allelic exchange vector pTOX5 cmR/msqR. ^73^ 800-1000 bp regions flanking the gene to be deleted were amplified from N16961 genomic DNA by PCR, cloned into the suicide vector pTOX5, and transformed into *E.coli* DH5α. The chloramphenicol-resistant colonies were screened for insertion of the flanking regions by colony PCR, and then verified using Sanger or whole-plasmid sequencing. The correct plasmid was transformed into *E.coli* SM10 or MFD. Conjugation was performed by spotting 15μL of the donor (*E.coli* SM10 or MFD) and 30μL of the recipient (*V. cholerae*) onto an LB plate containing 1% (wt/vol) glucose (+ 600μM DAP if using MFD as donor), and incubated for 4hrs at 37°C or overnight at 30°C. First selection was done by streaking the mating mixture out on LB plates containing chloramphenicol (5μg/mL), streptomycin (200μg/mL), and 1% glucose, and incubated overnight at 30°C. Chloramphenicol-resistant and streptomycin-resistant colonies were grown without selective pressure in LB broth containing 1% glucose for 4-6hrs at 37°C, followed by counter-selection done by streaking 2-4 candidates on M9 minimal medium plates supplemented with 0.2% casamino acids, 0.5mM MgSO_4_, 0.1mM CaCl_2_, 25μM FeCl_3_ in 50μM citric acid and 2% rhamnose. Single colonies were further streaked out on M9-streptomycin-rhamnose plates and verifired by colony PCR using flanking and internal primers for the gene of interest. Further verification was done by whole-genome sequencing in some cases.

Ectopic chromosomal expression from an IPTG-inducible *P_tac_* promoter was done using the suicide vector pTD101, a pJL1 derivate carrying the *P_tac_* promoter, *lacI^q^*, *sacB* gene and promoter, MCS (downstream of *P_tac_*), and allows integration of the gene of interest into the native *V. cholerae lacZ* (*vc_2338*) locus. ^74^ Genes of interest were amplified from N16961 genomic DNA by PCR, cloned into pTD101 introducing a strong RBS (AGGAGG) and spacer region upstream of the ORF. Selection was done on LB plates containing streptomycin (200μg/mL), carbenicillin (200μg/mL) and X-gal (120μg/mL) followed by counter-selection on salt-free LB agar plates containing 10% sucrose, streptomycin (200μg/mL) and X-gal (120μg/mL).

Ectopic plasmid expression from an IPTG-inducible *P_tac_*promoter was done using the vector pHL100mob, a conjugatable derivate of pHL100. For depletion of genes, the suicide vector pATdep1 was used to place the gene of interest under an aTc-inducible promoter. Cloning was done as mentioned above, and selection of transconjugants was done on LB agar plates containing streptomycin (200μg/mL) and kanamycin (50μg/mL) for pHL100 or chloramphenicol (5μg/mL) for pATdep1 integration.

The *lacZ* transcriptional reporters were built by amplifying the desired promoter region (600bp upstream of *zapC*) and cloning into the suicide vector pAW61, a pJL1 derivate. pAW61 carries MCS, *E.coli* lacZ ORF, homology to *V. cholerae*’s *lacZ* locus (like pTD101), allowing replacement of the native *lacZ* with the promoter-*lacZ* fusion. Strains were generated as described for pTD101 except that X-gal was not used, and the right insertions were verified using colony PCR using primers annealing to *zapC* promoter region and downstream flanking *V. cholerae*’s *lacZ* region.

### TnSeq and analysis

TnSeq was performed as described previously ^75,76^. WT and Δ6LTG strains (3 replicates each) were mated with *E. coli* MFD λ pir harboring the pSC189 suicide plasmid encoding the mariner transposon. The transconjugants were selected on LB agar plates containing Kanamycin 50 μg/mL to select for transposon insertions and approaximately 200,000 colonies/mating were obtained. The colonies were resuspended in 30mL LB, 1/5^th^ of which was used for genomic DNA (gDNA) extraction, and the rest was frozen in 30% glycerol as stock in −80°C. The gDNA was then fragmented using NEB fragmentase, followed by blunting (blunting enzyme mix; NEB), A-tailing, ligation of specific adaptors, and PCR amplification of transposon-DNA junctions using transposon and adaptor-specific primers. Libraries were sequenced on Illumina MiSeq. ^70^

Data analysis was performed using the MATLAB-based pipeline ARTIST.^76^ Genomic loci were predicted to be conditionally essential or enriched using the genome browser Artemis^77^, and insertion plots were generated in Graphpad prism.

### RNASeq and analysis

WT, Δ6LTG and Δ*dsvR* Δ6LTG strains (3 replicates each) were grown overnight in LB shaking at 30°C, diluted 1:100 into LB and grown shaking at 37°C until they reach mid-log phase (O.D._600_ ∼ 0.5). Cells were harvested and RNA was extracted using mirVana miRNA Isolation Kit (Invitrogen, AM1560). Genomic DNA contamination removal, library preparation, and Illumina sequencing were performed by Transcriptional Regulation and Expression Facility (TREx) at Cornell Institute of Biotechnology.

The sequencing fastq files along with the N16961 genome (.fa.gz) and genome annotations (.gtf.gz) were uploaded to galaxy ^78^. Adapter sequences were removed using trimmomatic, trimmed reads were aligned to genome using HISAT2, read counts per gene were quantified using htseq-count, and a raw counts (.txt) was generated for each pair-wise comparison. The differentially expressed genes were then identified using the DESeq2 analysis script in R ^79,80^. Genes with fold-change>2 or fold-change<0.5, p-value < 0.05 were considered for further analysis. Heatmaps were generated using Graphad Prism.

### Microscopy

Cells were grown in liquid media as mentioned before, and imaged without fixation on agarose pads (containing LB +/− antibiotic + 0.8% agarose) using Leica DMi8 inverted microscope. For time-lapses, image frames were 5 mins apart, with the stage temperature set at 37°C using PECON TempController 2000-1.

### Image analysis

To analyse phase contrast images, they were segmented using Omnipose^81^ using the bact_phase_omni model, and the masks were exported as .png files. These masks were imported and analyzed using MicrobeJ plugin in ImageJ/Fiji using default parameter ^82,83^. Stastical tests were run and plots were made in Graphpad Prism. To determine significant differences, Welch’s two-sample t-test was used. To analyze time-lapses, OmniSegger was used ^84^ and resulting values were analyzed using custom R scripts or Graphpad Prism.

### Growth curves

Cultures were grown overnight in LB shaking at 30°C, and diluted 1:100, 1:1000 or 1:10,000 into 200μL LB medium in a 100-well honeycomb plate. The plate is incubated at 37°C, with shaking at maximum amplitude, and the optical density of each well is measured at 600nm (O.D._600_) every 10 minutes in a Bioscreen C plate reader (Growth curves America).

### Spot dilutions

*V. cholerae* cultures were grown overnight in LB shaking at 30°C, and diluted 1:100 into 5mL LB medium and grown to exponential phase. Overnight or exponential cultures were 10-fold serially diluted in a 96-well plate until 1:10^7^, finally 5μL of all the dilutions 10^0^ through 10^7^ were spotted on LB agar plates (+/− antibiotics, +/− inducers). Plates were incubated overnight at 37°C.

### β-galactosidase assay

*V. cholerae* strains containing transcriptional-*lacZ* reporter constructs were grown overnight in LB and diluted 1:100 into LB, grown shaking at 37°C. Mid-log phase cultures (O.D._600_ ∼ 0.5) were harvested, and the β-galactosidase activity was quantified using o-nitrophenyl-β-galactoside as the substrate. ^85,86^

## SUPPORTING INFORMATION

Supplemental File 1: TnSeq analysis file

Supplemental File 2: RNASeq analysis file

Supplemental File 3: Strains, plasmids, and primers

## ACKNOWLEDGEMENTS

The authors thank Dr. Misha Kazi for assistance with the RNASeq analysis pipeline. We also thank the Cornell Statistics Consulting Unit, and in particular Dr. Lynn Johnson, for assistance with R scripts.

